# Somatic and terminal CB1 receptors are differentially coupled to voltage-gated sodium channels in neocortical neurons

**DOI:** 10.1101/2022.08.12.503665

**Authors:** Luke Steiger, Timur Tsintsadze, Glynis B. Mattheisen, Stephen M. Smith

## Abstract

Endogenous cannabinoid signaling is vital for important brain functions and can be modified pharmacologically to treat pain, epilepsy, and posttraumatic stress disorder. Endocannabinoid mediated changes to excitability are predominantly attributed to 2-arachidonoylglycerol at synapses. Here we identify a pathway in the neocortex by which anandamide, the other major endocannabinoid, powerfully inhibits sodium conductances in the soma resulting in a loss of neuronal excitability. This pathway is mediated by the cannabinoid receptor, and its activation results in a decrease of recurrent action potential generation. The synthetic cannabinoid, WIN 55,212-2, also inhibits VGSC currents indicating this pathway is positioned to mediate the actions of exogenous cannabinoids.

**Highlights:** Anandamide (AEA), a major endocannabinoid, indirectly inhibits VGSC currents in neocortical neurons.

This prevalent signaling pathway involves AEA activation of CB1 and other G-protein-coupled receptors localized to the intracellular compartment of neurons.

CB1 activation by AEA reduces VGSC availability at the soma but not at the axonal compartment suggesting tighter functional coupling between VGSCs and CB1 at the cell body.

Cannabinoid action on somatic CB1 inhibits VGSCs with high efficacy, providing a parallel pathway outside of nerve terminals, by which these ligands reduce neuronal excitability in the neocortex.

## Introduction

Endocannabinoids (eCBs) strongly regulate neuronal excitability in the central nervous system (CNS) and operate primarily by decreasing synaptic activity via the canonical cannabinoid receptor (CB1). CB1, the most abundant G-protein coupled receptor in the brain (Marsicano and Lutz, 1999), is concentrated at nerve terminals (Fitzgerald et al., 2013; Freund et al., 2003). Anandamide (AEA), the first discovered eCB (Devane et al., 1992) has a powerful influence on important neurobiological mechanisms. Perturbance of AEA metabolism results in profound changes to activity levels, temperature regulation, nociception, and sleep in animals and humans (Cravatt et al., 2001; Habib et al., 2019). Furthermore, AEA levels regulate fear extinction and recovery from posttraumatic stress disorder (Marsicano et al., 2002; Mayo et al., 2020). However brain AEA levels are far lower than those of 2-arachidonoylglycerol (2-AG), the other major eCB (Stella et al., 1997). Moreover 2-AG depresses synaptic transmission at a majority of synapses by acting via CB1 as a retrograde messenger (Tanimura et al., 2010; Yoshino et al., 2011). At a subpopulation of hippocampal synapses, AEA enhances synaptic transmission (Kim and Alger, 2010) and also strengthens inhibitory transmission via TRPV1 (Chávez et al., 2010; Chávez et al., 2014). However, these actions of AEA at a subgroup of hippocampal synapses are inadequate to account for the wide-ranging actions of AEA on behavior.

AEA has been reported to inhibit voltage-gated sodium channels (VGSCs) directly in a number of preparations (Al Kury et al., 2014; Kim et al., 2005; Nicholson et al., 2003; Okura et al., 2014). Since VGSC activation is essential for action potential generation, AEA regulation of VGSCs represents an alternative mechanism by which endocannabinoids regulate neuronal excitability. However, a number of key questions remain unanswered about this action in the CNS. What are the relative contributions of AEA and 2-AG on the regulation of VGSC? Are all compartments of highly polarized neurons equally sensitive? What is the mechanism by which AEA influences VGSC activity, how strong is this effect, and what neurons are affected? Here we address these questions by directly examining VGSC currents in the soma, dendrites, and boutons of neocortical neurons. We show that the majority of neocortical neurons are sensitive to AEA, that AEA causes stabilization of the inactivated channel state, and that AEA blocks VGSC currents indirectly at the soma. CB1 is a key effector at the soma and functional and structural data indicate intracellular CB1 receptors probably mediate this effect. In contrast, CB1 plays no role in regulating VGSC currents at the nerve terminal. The abundance and strength of VGSC inhibition by AEA via CB1 indicates this pathway is positioned to modulate neuronal excitability physiologically and also explain the anti-epileptic and analgesic actions of cannabinoid agonists.

## Results

### Anandamide inhibits neocortical VGSC currents with high efficacy

AEA was reported to directly inhibit VGSC currents in excitable cells and heterologous expression systems (Al Kury *et al*., 2014; Okura *et al*., 2014). We examined the prevalence, efficacy, kinetics, and state dependence of VGSC current inhibition by AEA in voltage-clamped neocortical neurons expressing multiple VGSC α-subunits (Katz et al., 2018). VGSC currents, evoked by stepping the membrane voltage to -10 mV from -80 mV at 0.2 Hz, decreased following the application of 10 µM AEA (Figure 1B,C). The response to AEA was observed in the majority of whole-cell somatic recordings (97%, n = 218) and the amount and rate of block of VGSC currents increased at depolarized holding potentials (V_h,_ Figure 1C). VGSC current inhibition by AEA was also concentration-dependent (Supplementary Figure 1). Direct inhibition by other sodium channel inhibitors (SCIs) is usually well described by a simple exponential time course (Jo and Bean, 2017). Unexpectedly, the time course of AEA inhibition of VGSC currents (I(t)), was better described by equation (1):

**Figure 1.**
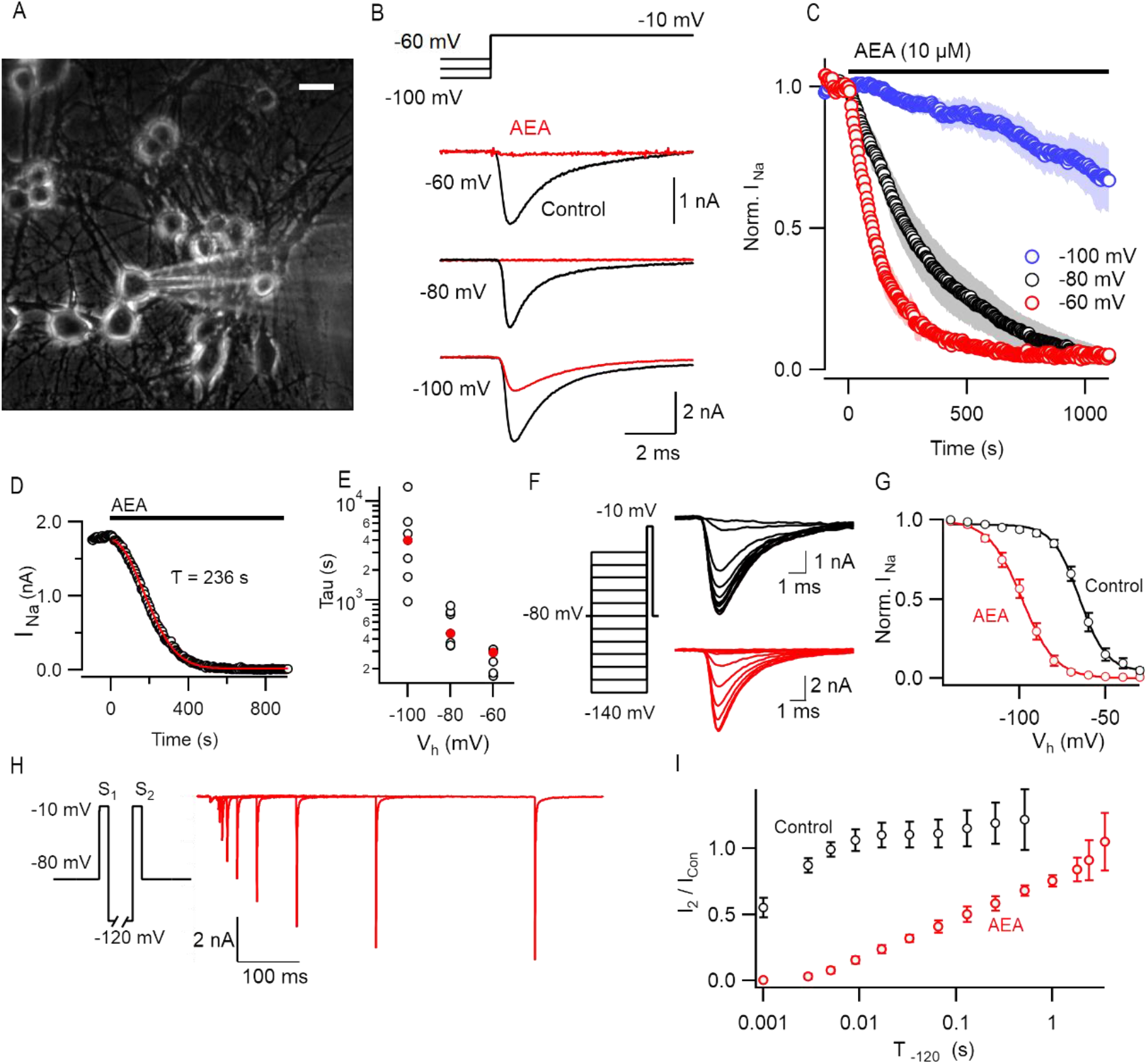
VGSC current block by anandamide increases with depolarization of the holding potential. A) Photomicrograph of primary cultured murine neocortical neurons White horizontal bar respresents 25 μm. B) Exemplar VGSC currents before (black) and after AEA (red; 10 μM for 400 seconds) application at three holding potentials. Neocortical neurons were stepped to -10 mV for 30 ms from a holding potential (V_h_) of -60, -80, or -100 mV . C) Normalized VGSC current following AEA application at time zero (indicated by black horizontal bar here and in later Figures). Data are shown as means ± S.E.M. with V_h_ represented as red (−60 mV, n=7), black (−80 mV, n=7), or blue (−100 mV, n=7). D) The time course of response to AEA in exemplar recording, shows that VGSC current amplitude is well described by equation 1. E) Time constants (r) of inhibition of VGSC currents by AEA is dependent on V_h_. F) VGSC currents activated following 100 ms conditioning prepulses in two neurons before (black) and after AEA (red) application. Voltage traces shown as left inset. G) Average normalized conductance plots following inactivating 100 ms prepulse indicates that AEA (red, n=8) shifts V_0.5_ by -33 mV (−65 vs -98 mV) compared to control (black, n=6). The curves represent the Boltzmann equation. H) Superimposed current traces (red) showing the time dependence of recovery of AEA-mediated VGSC inhibition following a voltage step to -120 mV (black). The current elicited by S_2_ (I_2_) increases with time. Voltage protocol shown in inset indicates step (S_1_ and S_2_) to -10 mV separated by step to -120 mV. I) I_2_ increased relative to the control VGSC current (I_Con_) elicited by the step from -80 mV to -10 mV before AEA application. The rate of recovery of I_2_ / I_Con_ is slowed substantially by AEA (red, n=4) compared to control (black, n=4).

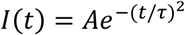

where A, t, and *τ* represent a constant, time, and the time constant respectively (Figure 1D). The *τ* decreased 14-fold as the holding potential was depolarized over a 40 mV range (Kruskal-Wallis test, P = 0.0004, Figure 1E) consistent with AEA preferentially inhibiting an inactivated channel state (Jo and Bean, 2017). Preferential binding of an antagonist to a specific channel state will impact the dynamic equilibrium altering the relative fraction of the channels that exist in a particular state and consequently the voltage-dependence of gating (Bean et al., 1983; Hille, 1977; Hille, 1978). We tested if VGSC inactivation was shifted by AEA using a 100 ms conditioning prepulse (Figure 1F,G). AEA affected VGSC gating, shifting the average half-maximal voltage (V_0.5_) by -33 mV, which is consistent with stabilization of the inactivated state. We tested if prolonged hyperpolarization reversed AEA inhibition of VGSC currents as observed for other direct and indirect SCIs (Mattheisen et al., 2018; Theile et al., 2016). VGSC currents (I_1_ and I_2_) were elicited by two 10 ms steps (S_1_ and S_2_) to -10 mV separated by a variable duration at -120 mV. In control experiments, I_2_ recovered to 50% of I_1_ within 0.8 ms. In contrast, after full block by AEA, I_2_ recovery was substantially slowed (130 ms for I_2_ to reach 50% of the control I_1_ value, Figure 1 H&I). Taken together, these data indicate that AEA strongly inhibits VGSC currents in the vast majority of neocortical neurons by stabilizing an inactivated channel state.

### Somatic VGSC Inhibition by AEA is mediated by CB1 receptor, and fully reliant on G-proteins

CB1 has been identified as the key receptor for endocannabinoid signaling in the CNS and is the most prevalent GPCR in the brain (Marsicano and Lutz, 1999). While 2-AG, has been shown to operate via CB1 at the synapse to impact neuronal excitability (Tanimura et al., 2010), AEA, phyto-, and synthetic cannabinoids have been proposed to inhibit VGSCs directly (Nicholson *et al*., 2003; Zhang and Bean, 2021). We re-evaluated the mechanism of action of AEA on VGSC currents because the time course of inhibition (Figure 1) appeared more consistent with an indirect effect (Mattheisen *et al*., 2018). Using a double pulse protocol (Figure 2A), we assayed the fraction of VGSC currents (I_1_ / I_2_) that were insensitive to AEA (10 µM). The first voltage step to -10 mV elicited the VGSC current resistant to AEA (I_1_) and the second step elicited the VGSC current (I_2_) after the majority of inhibition was reversed by a 1 s step to -120 mV (see Figure 1I). We observed a large range of values of I_1_ / I_2_ in CB1 null mutant neurons (*Cnr*^*-/-*^) and the neurons from wild-type litter mates (*Cnr1*^*+/+*^; Figure 2C) but block did not increase with the duration of incubation (within the range 20-100 min.). Deletion of CB1 resulted in an increase of the median value of I_1_/ I_2_ (P < 0.0001) from 0.50 (n = 21) to 0.85 (n = 31, Figure 2C) reflecting a substantial loss of sensitivity to AEA. Next, we tested if the time course of inhibition of VGSC currents by AEA was affected by CB1 deletion. Using a similar protocol as Figure 1B, voltage-clamped neurons were stepped from -80 mV to -10 mV at 0.2 Hz. AEA only reduced the VGSC currents in the *Cnr1*^*-/-*^ neurons by 18% compared with 65% in wild-type neurons (Figure 2D-G; n = 10 each, P < 0.001) indicating CB1 mediated the majority of the effect of AEA on VGSC currents. The time course of inhibition was also slowed in the *Cnr1*^*-/-*^ compared to *Cnr1* ^*+/+*^ neurons (Figure 2G; *τ* = 499 ± 96 s and 1639 ± 235 s, P = 0.0002).

**Figure 2:**
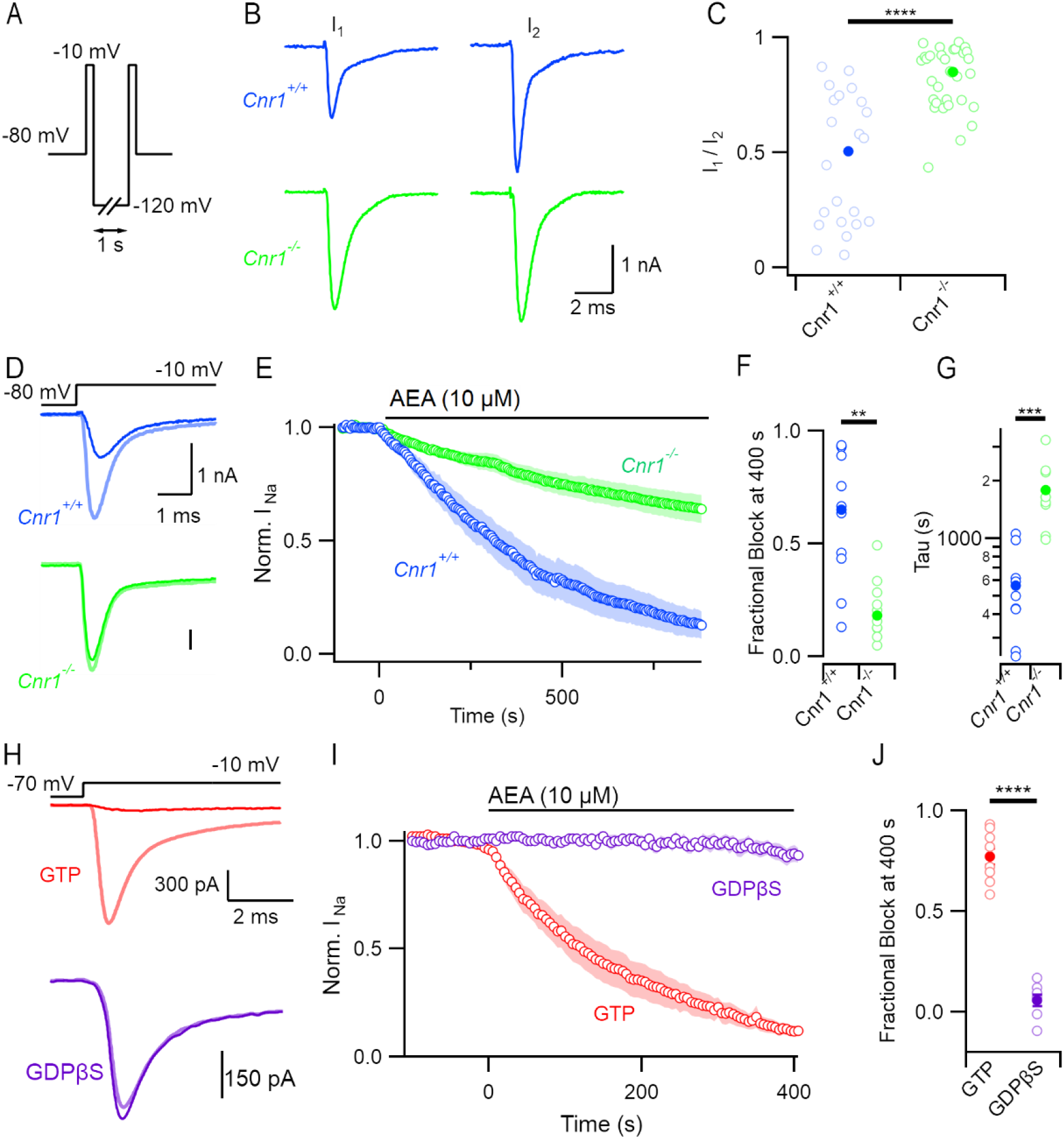
AEA block of VGSC is mediated by CB1.. A) Double-pulse voltage protocol illustrating two steps to -10 mV with hyperpolarizing period of 1 second at -120 mV to reverse inactivation before the second step. B) Representative double-pulse VGSC current traces from *Cnr1*^+/+^ (wild-type, blue) and *Cnr1*^-/-^ neurons (green) following incubation in 10 μM AEA. C) Ratio of VGSC amplitude (I_1_ / I_2_) from double pulse protocol in wild-type (median 0.50, n=21) and *Cnr1*^-/-^ (0.85, n = 31) neurons, Kruskal-Wallis (KW) test P <0.00001. Here and in later Figures P values designated as follows: * P < 0.05, ** P< 0.01, *** P< 0.001, and **** P< 0.0001. D) VGSC current traces before (light) and after (dark) 400 seconds of 10 μM AEA perfusion in *Cnr1*^+/+^ (blue) and *Cnr1*^-/-^ neurons (green). E) Time course of normalized VGSC current amplitude in *Cnr1*^+/+^and *Cnr1*^-/-^ neurons (n = 10 each) before and following application of 10 μM AEA. Cells were stepped from -80 mV to -10 mV for 30 ms every 5 seconds. F) Fractional block (1 - normalized residual current) at 400 seconds following exposure to AEA in *Cnr1*^+/+^ and *Cnr1*^-/-^ neurons; KW test P = 0.001. G) Time constants of inhibition as described by equation 1 in *Cnr1*^+/+^ and *Cnr1*^-/-^ neurons; KW test P = 0.001. H) Exemplar VGSC current traces before and 200 seconds after application of AEA with (purple, n = 6) or without (red n = 8) 2mM GDPβS. Neocortical neurons were stepped from a V_h_ of -70 to -10 mV every 5 seconds. I) Normalized VGSC current amplitude prior to and following application of 10 μM AEA alone or in the presence of GDPβS (2mM) in the recording pipette. J) Fractional block of VGSC following 400s of 10 μM AEA application with or without GDPβS. The blocked current fraction was 0.06 ± 0.03 vs. 0.77 ± 0.07, n = 6 and 8 respectively; P < 0.0001.

To confirm that AEA-induced block of VGSC currents involved G-protein signaling, we utilized GDPβS, which affects G-protein cycling by inhibiting GTP binding (Eckstein et al., 1979; Suh et al., 2004). AEA (10 µM) inhibited VGSC currents by 77 ± 7% (n = 8), 400 seconds after the onset of application in these recordings (V_h_ was -70 mV). In contrast, inclusion of GDPβS (2 mM) in the recording pipette substantially attenuated the AEA effect (Figure H-J; 6 ± 3%, n = 6, P < 0.0001) indicating that any direct inhibition by AEA represents a minor component of its overall action on VGSC currents.

Next, we tested if CB1 antagonists affected the inhibition of VGSC currents during the application of AEA. The maximum fractional block of VGSC currents by AEA was reduced from 0.96 (n = 7) to 0.27 (n = 6) by the neutral CB1 antagonist AM4113 (Figure 3A,B; 5 µM plus DMSO versus vehicle in pipette solution; P = 0.0008). Likewise, the CB1 inverse agonist AM251 (5 µM in pipette solution) reduced the maximum fractional block of VGSC currents by AEA from 0.96 (n = 7) to 0.45 (Figure 3C,D; n = 4; P = 0.014). AM4113 was ineffective in the *Cnr1*^*-/-*^ neurons, indicating that the antagonist was acting at CB1 to prevent AEA inhibition of VGSC currents (Figure 3E,F; P = 0.58).

**Figure 3:**
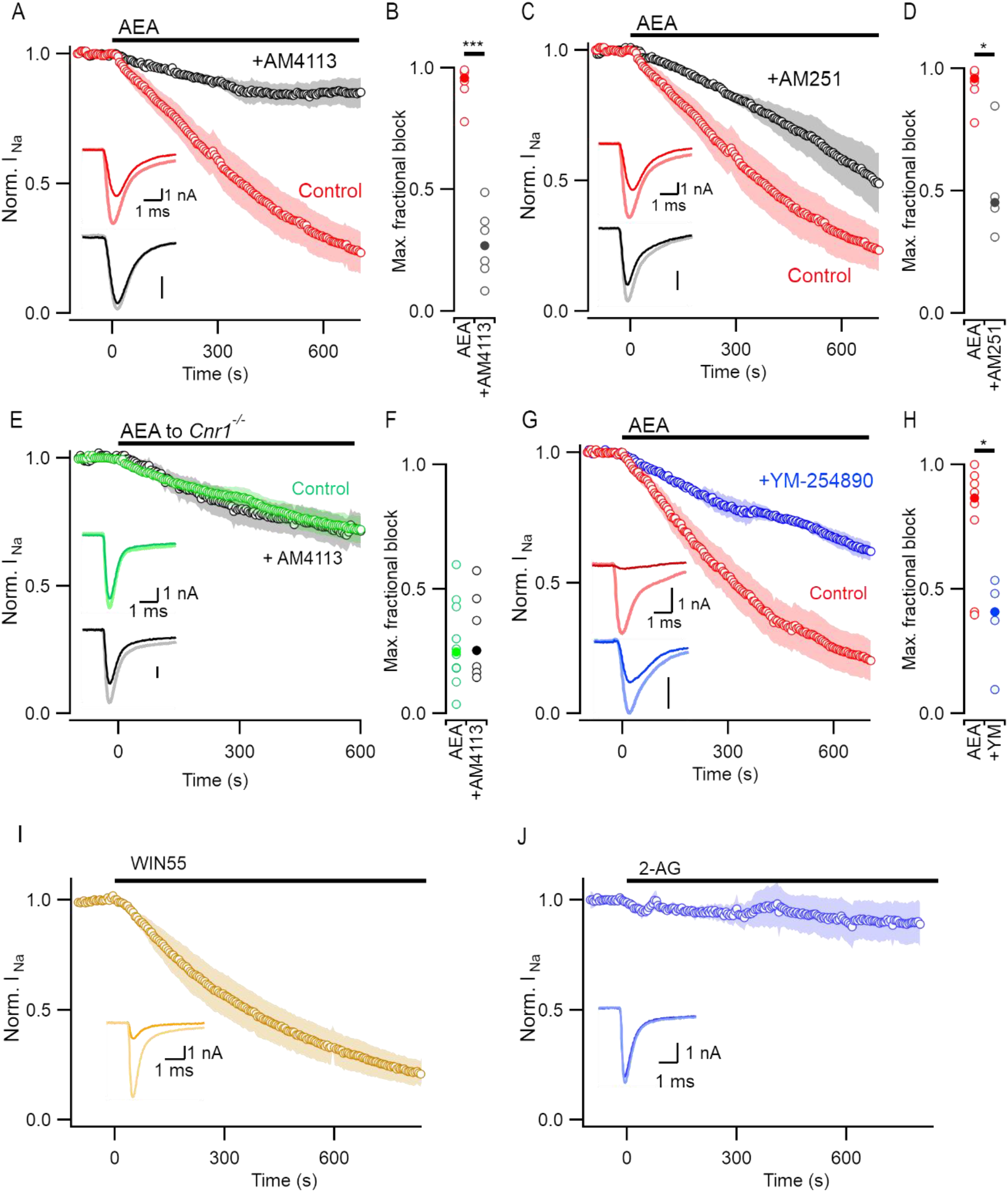
Intracellular CB1 antagonists attenuate block by AEA. A) Time course of VGSC current inhibition by AEA (plus DMSO, red, n = 7) versus AEA + 5 µM of the neutral CB1 antagonist AM4113 (black, n = 6). VGSC current traces represent application before (light) and after (bold) application of 10 µM AEA. Neurons were stepped to -10 from a Vh of -80 mV every 5 seconds throughout. B) Maximal fractional VGSC current block observed was reduced from a median 0.96 to 0.27 in the presence of AM4113 (black) vs equivalent DMSO vehicle control (red); P = 0.0008. C) Time course of VGSC current inhibition by AEA (plus DMSO, red, n = 7) versus AEA + 5 µM of the inverse CB1 agonist AM251 (black, n = 4). VGSC current traces represent application before (light) and after (bold) application of 10 µM AEA. Neurons were stepped to -10 from a Vh of -80 mV every 5 seconds throughout. D) Maximal fractional VGSC current block observed was reduced from a median 0.96 to 0.45 in the presence of AM251 (black) vs equivalent DMSO vehicle control (red); P = 0.014. E) Time course of VGSC current inhibition by AEA+AM4113 [See A)] applied to *Cnr1*^-/-^ neurons (black, n = 5) versus AEA alone (green, n = 10). VGSC current traces represent application before (light) and after (bold) application. F) Maximal fractional VGSC current block observed by application of AEA or AEA+AM4113 to *Cnr1*^-/-^ neurons (median 0.248 vs 0.253; P = 0.57). G) Time course of VGSC current inhibition by AEA/DMSO (red, n = 7) versus AEA + 500 nM YM-254890 (blue, n = 5). VGSC current traces represent application before (light) and after (bold) application of 10 µM AEA. H) Maximal fractional VGSC current block observed was reduced from a median 0.86 to 0.41 in the presence of YM-254890 (blue) vs DMSO vehicle control (red); P = 0.048. I) Time course of VGSC current before and during perfusion of the synthetic CB1 agonist Win-55 212-2 (n = 4). VGSC current traces represent application before (light) and after (bold) application of 10 µM Win-55. J) Time course of VGSC current before and during perfusion of the endogenous cannabinoid 2-arachidonoylglycerol (2-AG, n = 4). VGSC current traces represent application before (light) and after (bold) application of 10 µM 2-AG.

The partial sensitivity of *Cnr1*^*-/-*^ neurons to AEA (Figure 2C,F), the incomplete block of AEA by AM4113 (Figure 3D) and the effectiveness of GDPβS indicate that another GPCR may be combining with CB1 to transduce the effect of AEA. We tested the sensitivity of VGSC currents to other cannabinoid agonists to characterize the pharmacological characteristics of the activated CB1 receptor. The synthetic CB1 agonist WIN 55,212-2 (10 µM) strongly reduced VGSC current amplitude (Figure 3I; n = 3) whereas 2-AG (10 µM), another endocannabinoid, was ineffective (Figure 3J, n = 4).

The signaling pathway between CB1 and the VGSCs is unclear since CB1 signals via a number of G-protein subtypes (Diez-Alarcia et al., 2016). YM-254890, the small molecule G_q/11_ inhibitor (500 nM, (Uemura et al., 2006)) reduced maximum AEA-mediated block of VGSC currents from 86% (n = 8) to 41% (Figure 3G,H; n = 5; P = 0.049). This substantial attenuation of the action of AEA suggests a role for G_q/11,_ however YM-254890 may also inhibit other G-proteins (Peng et al., 2021). Taken together, our experiments using *Cnr1*^*-/-*^ neurons, CB1 antagonists, and inhibitors of G-proteins indicate that CB1 is responsible for the majority of inhibition of VGSC currents by AEA and that this is mediated via a GTP-dependent mechanism.

### Subcellular location and function of CB1

The utilization of photolysis to activate signaling lipids that are restricted to intracellular or extracellular compartments has illustrated the functional importance of the subcellular localization of cannabinoid receptors in excitable cells (Frank et al., 2018; Nadler et al., 2015). CB1 ligands are lipophilic and it has been assumed that these agents penetrate tissue uniformly. We tested if AM4113 (5 µM) affected AEA inhibition of VGSC currents when applied extracellularly and preincubated before AEA application (20 min.) to ensure its effectiveness. The median I_1_ / I_2_ ratio after AEA incubation (10 µM for 20-90 min.) was not significantly changed by AM4113 (Figure 4A,B; 0.53 to 0.69, P = 0.32) indicating reduced CB1 antagonist effectiveness when applied externally (see Figure 3A,B). Consistent with this finding, AM251 (5 µM) did not alter the inhibition of VGSC currents by AEA when applied externally (Figure 4C).

**Figure 4.**
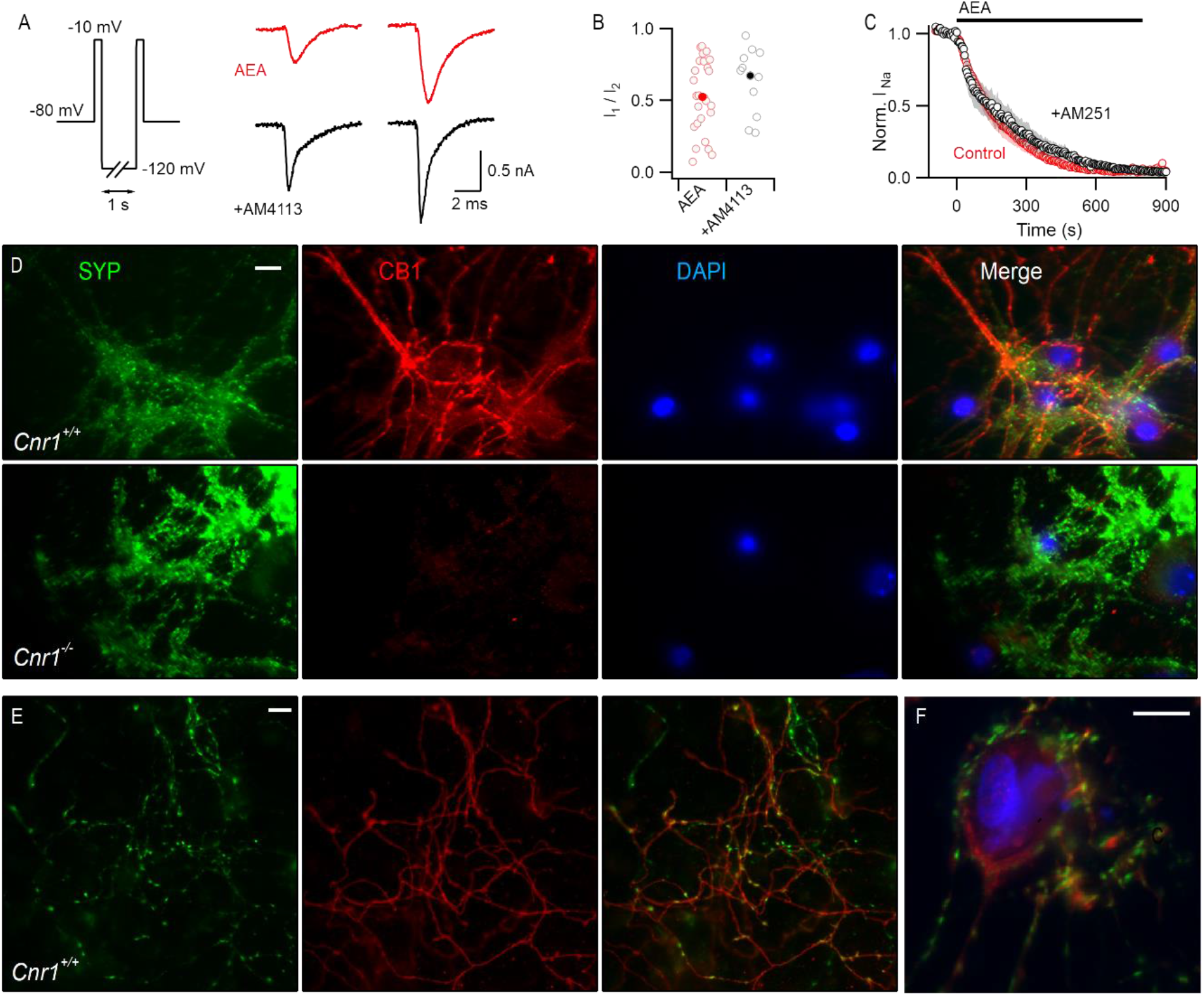
Extracellular antagonists do not attenuate AEA block of VGSC; CB1 receptor is expressed on axons and within somata of neocortical neurons. A) Exemplar currents elicited by double-pulse protocol after incubation in AEA (10 µM) or AEA plus AM4113 (5 µM). B) Ratio of VGSC amplitude (I_1_ / I_2_) from double pulse protocol in AEA (red, n = 26) or AEA +AM4113 (black, n = 13), median 0.53 vs 0.69; P = 0.32. C) Normalized VGSC current amplitude prior to and during perfusion of AEA (red) or AEA + 10 µM AM251 (black). Neurons were stepped to -10 from a V_h_ of -70 mV. D) Fluorescent microscopy image of antibody-labeled sister *Cnr1*^+/+^ and *Cnr1*^-/-^ neocortical neuron cultures. Cells were stained for synaptophysin 1 (SYP, green), cannabinoid 1 receptor (CB1, red) and nuclei (DAPI, blue). Images capture under 60x magnification. Scale bar 10 µm. E) Fluorescent microscopy image of antibody-labeled synaptophysin (green) and CB1 (red) on neocortical neuron processes. Images captured under 60x magnification. Scale bar 10 µm. F) Fluorescent microscopy image of neocortical neuronal cell body depicting synaptophysin 1 (green), CB1 (red) and DNA (blue). Images taken under 60x magnification. Scale bar 10 µm.

The enhanced effectiveness of CB1 antagonists applied via the pipette solution may indicate a role for intracellular CB1 in the AEA-mediated inhibition of VGSC currents. Fixed neocortical neurons were imaged to examine the distribution of CB1. Using synaptophysin as a presynaptic marker we co-stained for CB1 and nuclei (DAPI). The most intense CB1 staining was observed in the neuronal processes of *Cnr1*^*+/+*^ neurons (Figure 4C). On less dense areas of the cultures, synaptophysin was clearly concentrated at puncta, reflecting boutons, with less intense staining of the processes (Figure 4D). In contrast, CB1 co-stained a fraction of the puncta while clearly delineating the interconnecting axons, indicating localization of CB1 to axons and boutons (Figure 4D). By focusing on nuclei identified with DAPI, we detected CB1 staining throughout the neuronal soma in *Cnr1*^*+/+*^ cultures reflecting a distribution consistent with an intracellular localization (Figure 4E) as noted by others (Fletcher-Jones et al., 2020).

CB1 plays an important role at nerve terminals by reducing release probability (Tanimura *et al*., 2010). We asked if the axonal CB1 (Figure 4D) also contributed to VGSC inhibition by AEA. Using a modified patch-clamp method (Ritzau-Jost et al., 2021) we recorded VGSC currents directly from small neocortical boutons and the associated axon. We identified boutons by extracellular loading with the styryl dye, FM1-43 (Smith et al., 2004), and confirmed correct patch pipette placement by including a complementary dye (Atto 594) in the pipette solution (Figure 5A). VGSC currents were elicited with a step from -80 mV to -10 mV at 0.2 Hz and AEA (10 µM) applied to the recording after a stable VGSC current was established. AEA (800 s) blocked the VGSC current by 58 ±10% (n = 6) and this action was unaffected by deletion of CB1 (Figure 5; 45 ±11%, n = 9; P = 0.52). Following the application of AEA we used the double pulse protocol to confirm that any AEA-mediated inhibition of VGSC currents was reversible (Figure 5C,F). The fraction of the VGSC current insensitive to AEA (I_1_ / I_2_ ratio) was consistent with the fractional block observed in the diary plots (Figure 5E,F). These data indicate that VGSC currents in axons and boutons are more resistant to AEA than the somatically recorded currents (Figure 1C). Furthermore, AEA acts independently of CB1 at the bouton and axon in contrast to the soma where CB1 activation by AEA strongly inhibits VGSC current amplitude.

**Figure 5.**
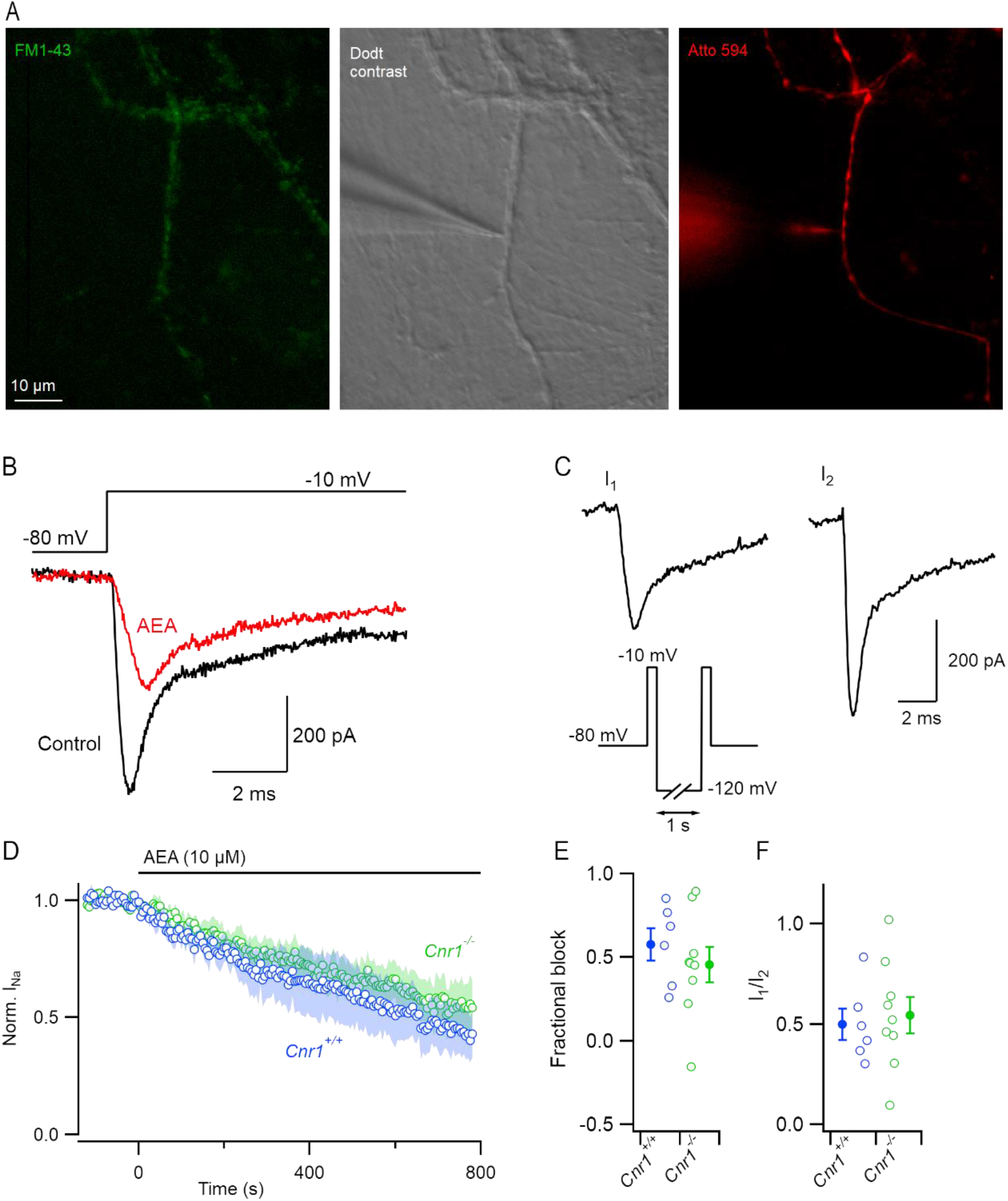
AEA inhibition of VGSC is modest at nerve terminals and not mediated by CB1 A) Photomicrograph of neocortical boutons. Boutons were identified via extracelluar FM1-43 (2mM, green), patch-clamped under dodt contrast (grey), and confirmed by inclusion of Atto 594 in the recording pipette (2mM, red). B) Voltage protocol and representative terminal VGSC traces before (red) and 800s after (black) perfusion of AEA. Currents were elicited with a voltage step from -80 to -10 mV. C) Double pulse protocol to -10 from either -80 (I_2_) or -120 (I_1_) mV used to facilitate recovery of VGSC current after block by AEA. D) Time course of normalized VGSC current amplitude before and during perfusion of AEA in wild-type (blue, n = 6) and *Cnr1*^-/-^ neurons (green, n = 9) E) Fractional block at 800 seconds following exposure to AEA in wild-type and *Cnr1*^-/-^ neurons, 0.58 vs 0.46 respectively; P = 0.52. Shaded points represent mean ± SEM F) Ratio of VGSC amplitude (I_1_ / I_2_) from double pulse protocol after 800 seconds of AEA perfusion in wild-type (0.50) and *Cnr1*^-/-^ neuronal boutons. Shaded points represent mean ± SEM for each group.

The absence of dendritic CB1 staining suggested CB1 would not regulate dendritic VGSC currents. However, it possible that CB1 signaling from the somatic compartment could propagate to and reduce dendritic VGSC currents. We tested this hypothesis by recording directly from dendrites (Figure 6A). Overall VGSC currents in dendrites were much less sensitive to AEA than soma and axons (Figure 6B). The VGSC currents were inhibited by 24±16 % and 10±12 % by 800 s application of AEA (10 µM) in *Cnr1*^*+/+*^ and *Cnr1*^*-/-*^ neurons, n = 5 and 8, respectively (Figure 6E). Consistent with this, the I_1_ / I_2_ ratio was close to unity. In contrast to our findings at the soma, these data indicate axonal and dendritic VGSC currents are much less sensitive to AEA and this modest action is independent of CB1.

**Figure 6.**
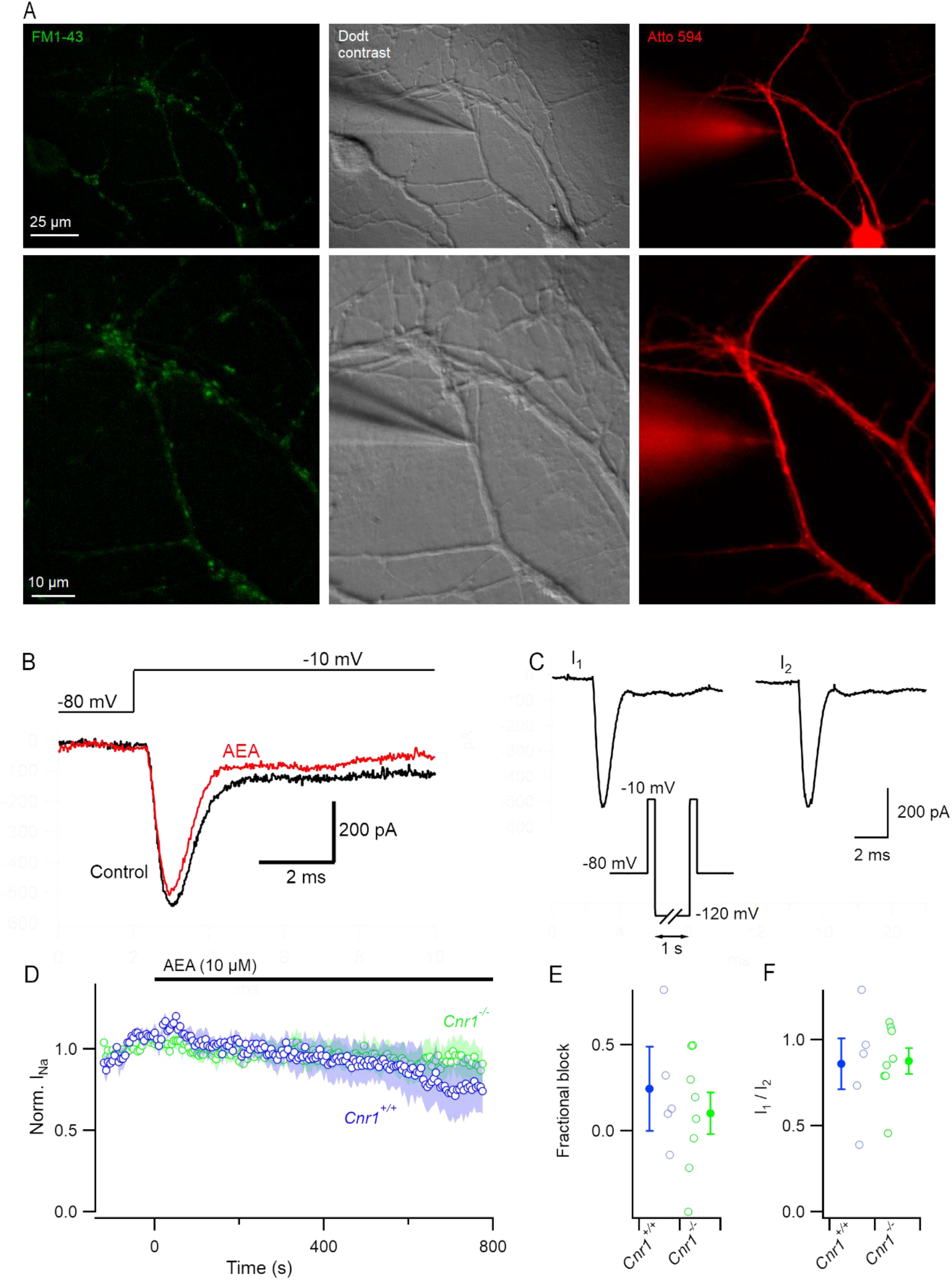
Dendritic VGSC are insensitive to AEA. A) Photomicrograph of neocortical dendrites. Boutons were identified via extracelluar FM1-43 (2mM, green), patch-clamped under dodt contrast (grey), and confirmed by inclusion of Atto 594 in the recording pipette (2mM, red). B) Voltage protocol and representative dendritic VGSC traces before (red) and 800s after (black) perfusion of AEA. Currents were elicited with a voltage step from -80 to -10 mV. C) Double pulse protocol to -10 from either -80 (I_1_) or -120 (I_2_) mV used to facilitate recovery of VGSC current after block by AEA. D) Time course of normalized VGSC current amplitude before and during perfusion of AEA in wild-type (blue, n = 5) and *Cnr1*_-/-_ neurons (green, n = 9) E) Fractional block of normalized dendritic VGSC current at 800 seconds following exposure to AEA in wild-type and *Cnr1*_-/-_ neurons, 0.24 vs 0.10 respectively. Shaded points represent mean ± SEM. F) Ratio of VGSC amplitude (I_1_ / I_2_) from double pulse protocol after 800 seconds of AEA perfusion in wild-type and *Cnr1*_-/-_ neuronal boutons. Shaded points represents mean ± SEM for each group.

### AEA block of VGSC is use-dependent, manifests as potent reduction in repetitive spiking

The impact of endocannabinoids on neocortical excitability is expected to be strong because of the high efficacy of AEA at the somatic CB1-VGSC pathway (Figure 1). Preferential binding to the inactivated VGSC states may enhance inhibition of VGSC currents by AEA during sustained periods of activity consistent with other SCIs. However, since AEA was less effective after simple incubation (Figure 2C) and because indirect SCIs may be relatively ineffective at duty cycles above 0.2 Hz (Mattheisen *et al*., 2018), we examined the use-dependence of AEA using a modified diary plot protocol to determine the fraction of block 200 s and 500 s after the onset of application (Figure 7A-C). AEA was substantially less effective if the VGSC currents were not elicited during the period of application. Re-initiation of the depolarizing steps 200 s after onset of application (blue squares) revealed that the fractional block of VGSC currents was substantially reduced (0.08 vs 0.44; P = 0.0058) compared with those recorded in control, continuously cycled neurons (red circles). Delaying the re-initiation of the voltage steps to 500 s after the onset of AEA application (black triangles), increased the difference in fractional block (Figure 7A-C; 0.21 vs 0.87; P = 0.00037). Accelerating the duty cycle to 1 Hz had minimal effect on VGSC inhibition by AEA suggesting the use-dependence is more pronounced at frequencies below 0.2 Hz (data not shown).

**Figure 7.**
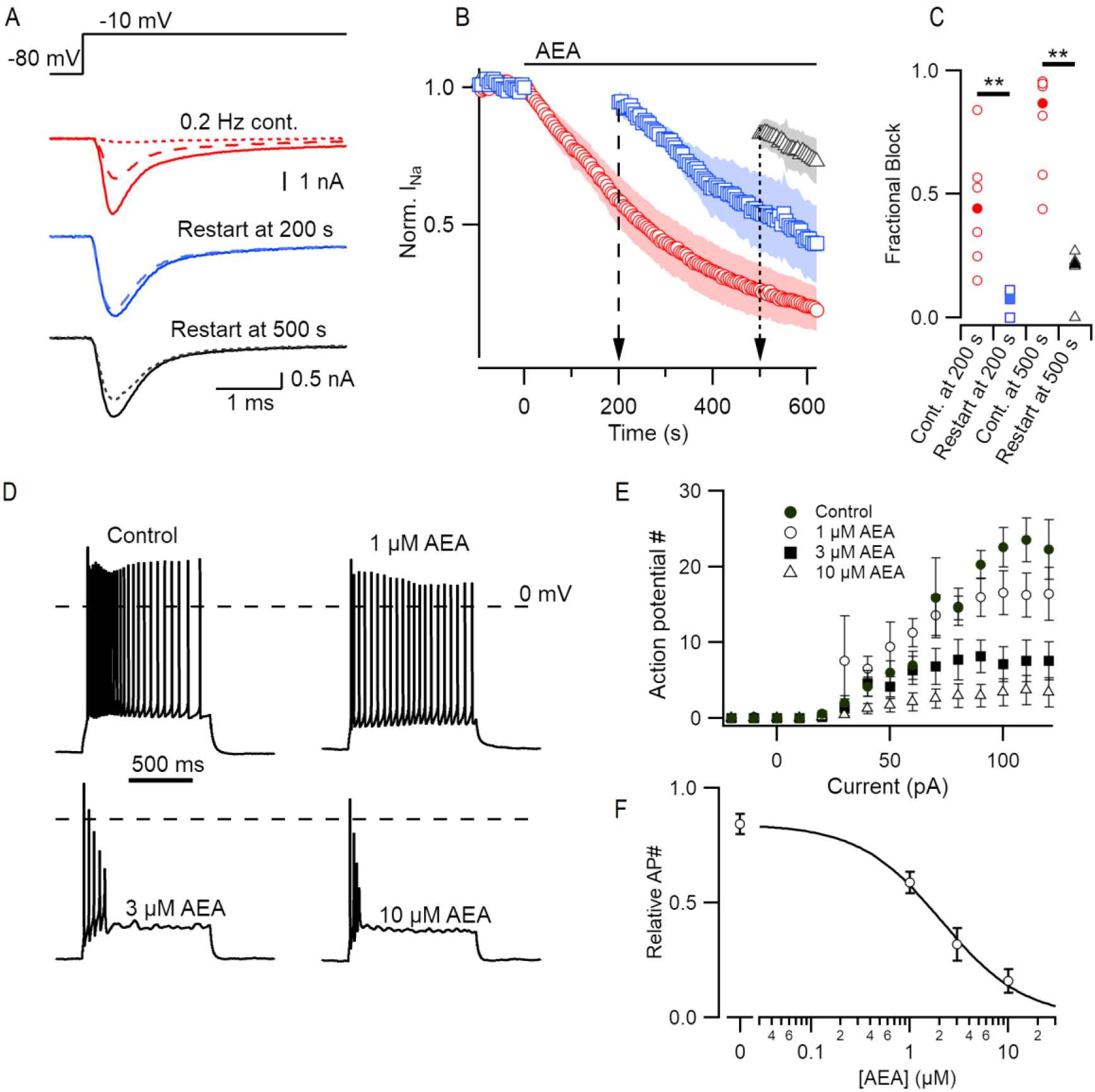
AEA inhibition of VGSC is use-dependent and results in reduced action potential generation in response to stepwise current injections. A) Exemplar VGSC traces of neocortical neurons continuously sampled at 0.2 Hz by voltage step from - 80 to -10 mV in the presence of AEA (red) versus those held silent for 200 (blue, dashed) or 500 (black, dotted) seconds. B) Time course of normalized VGSC current amplitude prior to and during application of AEA. Cells were sampled continuously every 5 seconds (0.2 Hz) or held inactively at -80 mV for either two (blue, circles) or five (black, triangles) hundred seconds before resuming. C) Fractional block of normalized VGSC current for cell sampled at 0.2 Hz (red dots) versus those held silent for 200 (blue squares, 0.08 vs 0.44; P = 0.0058) or 500 seconds (black triangles, 0.21 vs 0.87; P = 0.00037). D) Exemplar voltage traces within a single neuron firing in response to stepwise current injections after 500 seconds exposure to 1, 3, or 10 µM AEA. E) Number of action potentials generated in response to variable stepwise current injections (−20 to 120 pA) for each concentration of AEA. Error bars reflect SEM, n = 7 cells. F) The last 3 series of current injections (100-130 pA) were used to calculate a concentration-effect curve based on the normalized action potential number within each cell, EC50 = 2.1 μM AEA.

Next, we determined the concentration-effect relationship of AEA on intrinsic neocortical excitability. We measured the number of action potentials evoked by a series of 1 s current injections (0-120 pA; Figure 7) in neocortical neurons after synaptic transmission was inhibited with CNQX, APV and Gabazine (10, 50, and 10 µM respectively). AEA was applied for 500 s in increasing concentration thereafter, while V_h_ was held at -80 mV. Neuronal excitability was re-evaluated at 1, 3 and 10 µM AEA using the same series of current injections. Action potential number plateaued with the 100-120 pA injections and decreased as AEA was increased (Figure 7E). AEA reduced the number of action potentials generated in a concentration-dependent manner with a half-maximal concentration of 2.2 ± 0.1 µM (Figure 7F, n = 7 cells). AEA application reduced the peak action potential amplitude (82 ± 4 control vs. 62 ± 6 mV threshold to peak, P = 0.03, data not shown). Additionally, AEA application had little effect on the position of the first spike but reduced recurrent spikes (Figure 7D) suggesting that AEA will be more effective at active neuronal circuits.

## Discussion

Here we identify and characterize a mechanism by which AEA acts as a strong inhibitor of neuronal excitability at the cell body. Several features of this pathway are unexpected. First, contrary to prior reports, this is an indirect and GTP-dependent signaling pathway. Second, it is mediated by the CB1, the cannabinoid receptor which canonically operates primarily on the plasma membrane of synapses. Third, the pathway is localized at the soma and attributable to an intracellular population of CB1 receptors. Fourth, VGSC inhibition by AEA is state-dependent and exhibits an unusual form of use-dependence. Lastly, its high efficacy and prevalence combine to substantially reduce action potential generation at low-micromolar concentrations. The strength and ubiquity of this pathway provides a mechanism to explain how endo-, phyto-, and synthetic cannabinoids control neocortical excitability.

### CB1 is a key player in AEA inhibition of VGSC currents

Contrary to prior reports (Al Kury *et al*., 2014; Kim *et al*., 2005; Nicholson *et al*., 2003) we find that CB1 plays an important role in the inhibition of VGSC currents. Block by AEA was highly attenuated in neurons lacking a CB1 receptor (Figure 2). Antagonists for CB1 attenuated the action of AEA consistent with CB1 dependence (Figure 3). Like these other studies (Al Kury *et al*., 2014; Kim *et al*., 2005; Nicholson *et al*., 2003), we found that when applied extracellularly the CB1 antagonists were ineffective (Figure 4). The difference in efficacy of the two routes of application may at first seem surprising due to the lipid solubility of these agents, though other lipid signaling ligands have been shown to have different effects when released intra or extracellularly indicating the importance of relative concentrations across cellular compartments (Frank *et al*., 2018; Nadler *et al*., 2015). The effectiveness of GDPβS and G-alpha inhibitor, YM-254890, at inhibiting the action of AEA on VGSC currents, also support an indirect GPCR mediated action (Figures 2,3). YM-254890 inhibits G_q/11_ and G_s_ by preventing GDP exchange for GTP on these alpha subunits, and potentially also exhibits biased inhibition of G_i/o_ (Peng *et al*., 2021; Taniguchi et al., 2003). That YM-254890 attenuated but did not ablate AEA block, implies additional involvement of alpha subunits other than those sensitive to YM-254890. Saturation of the cytoplasm with GDPβS by applying it through the recording electrode completely abrogated block by AEA, strongly suggesting the pathway is entirely dependent on exchange of GDP for GTP. GDPβS acts similarly to YM-254890 by competitively inhibiting GTP binding to G-proteins, but is non-specific (Eckstein *et al*., 1979; Suh *et al*., 2004). This raises the interesting question of which G-protein subunits transduce the inhibitory signal of AEA. The cannabinoid receptor is promiscuous and couples with multiple G protein subtypes (Diez-Alarcia *et al*., 2016) based on cell type (Navarrete and Araque, 2010), subcellular location (Bénard et al., 2012), and dimerization with other receptors (Glass and Felder, 1997). In mouse cortex the synthetic AEA analog ACEA stimulates Gα_i1_, Gα_i3_, Gα_o_, and Gα_q/11_ via CB1 at relatively uniform levels (Diez-Alarcia *et al*., 2016). It seems likely that one or more of these subunits also mediates block of VGSC.

The partial block of VGSC currents in the *Cnr1*^*-/-*^ neurons could indicate a modest contribution of direct sodium conductance inhibition by AEA (Figure 2). However, the absence of effect of AEA in the presence of GDPβS suggests it is more likely that another GPCR is responsible. This interpretation is also consistent with the observation that AM4113 is incompletely effective. Since other GPCRs may heterodimerize with CB1 (Oyagawa and Grimsey, 2021), CB1 and the unidentified GPCR could be operating as homo- or heterodimer pairs to mediate AEA block of VGSC (Figure 8).

**Figure 8.**
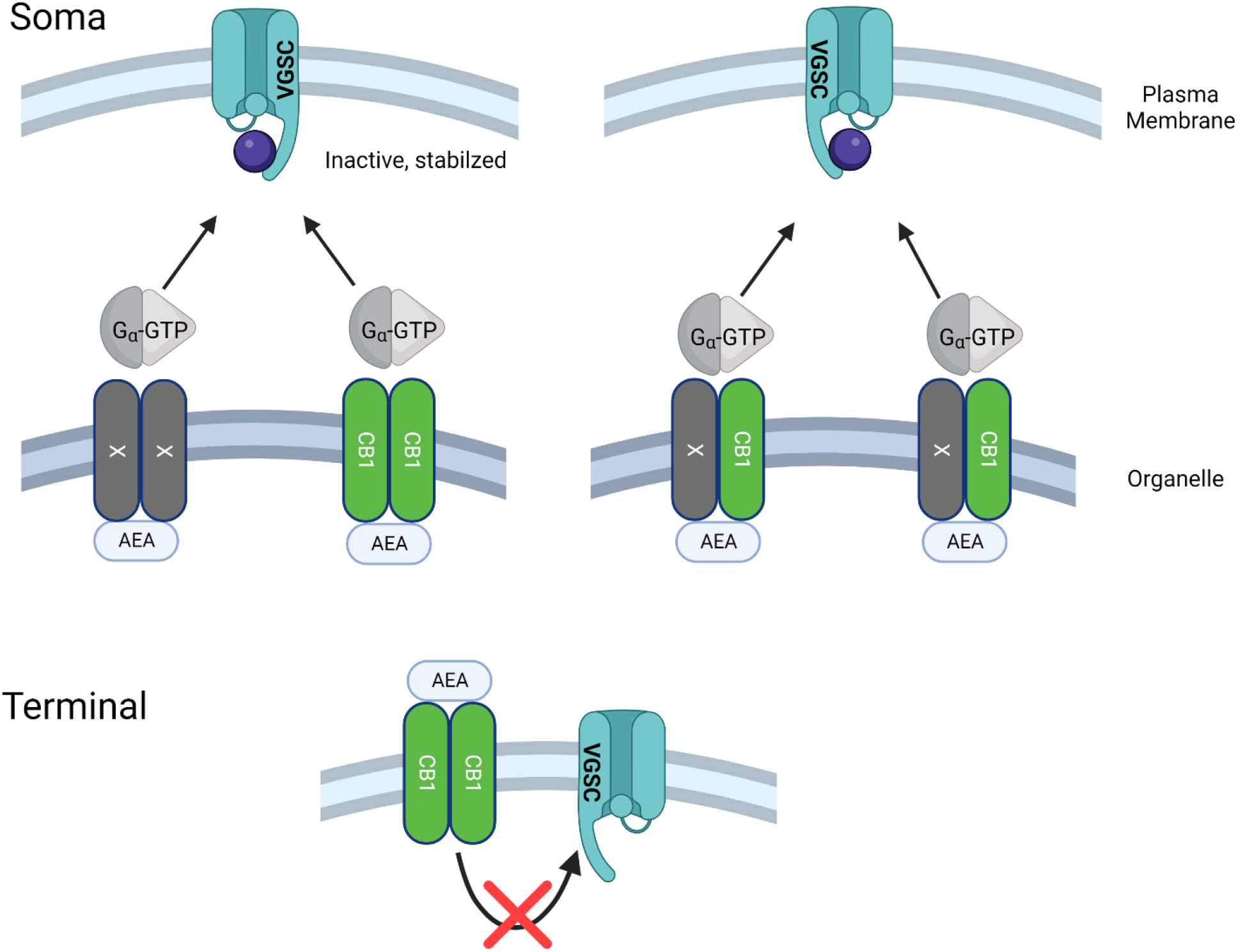
Cartoon depicting models for proposed signaling pathways by which AEA inhibits VGSCs. VGSCs in the plasma membrane of the soma are stabilized in the inactive state by G-proteins activated by AEA operating via GPCRs (CB1 (green) and CB1-independent (X, gray)). CB1 is shown localized to an intracellular organelle. In the left panel the GPCRs operate as homologous dimers and the non-CB1 component may be intracellular or on the plasma membrane. In the right panel, CB1 and X form heterologous dimers and are localized to an organelle. The lower panel shows that at the nerve bouton, CB1 does not transduce AEA signaling to VGSCs.

### The role of AEA

Endocannabinoid messengers have a well-established role regulating synaptic strength at many synapses in the hippocampus and neocortex which facilitates higher order processes such as learning and memory (Griebel et al., 2015; Marsicano and Lafenêtre, 2009). The most abundant synaptic endocannabinoid, 2-AG (Stella *et al*., 1997), is produced in the hippocampus as activity increases (Farrell et al., 2021) and binds to presynaptic CB1 receptors inhibiting adenylyl cyclase and calcium channels, thereby suppressing transmitter release (for review (Castillo et al., 2012)). Comparatively less is known about how AEA modulates excitability and behaviors. First characterized 30 years ago (Devane *et al*., 1992) AEA was rapidly established via behavioral studies as important for facilitating working memory and possessing powerful anxiolytic and analgesic properties (Mallet and Beninger, 1996; Terranova et al., 1995; Walker et al., 1999) including in humans (Habib *et al*., 2019). Here, we outline how AEA can regulate excitability by strongly inhibiting VGSC. The widespread reduction of VGSC currents in cortical neurons and high potency of AEA explain how this endocannabinoid can modulate neuronal excitability. By stabilizing the inactivated state of the VGSCs in neocortical neurons, AEA signaling reduces VGSC availability and decreases the probability of action potential generation (Milescu et al., 2010). This finding strongly augments the r effects of AEA on other ion channels, including TRPV1. The high prevalence of CB1 and VGSCs in neocortical neurons contrasts with TRPV1 (Cavanaugh et al., 2011; Marrone et al., 2017) suggesting that the majority of the effect of AEA on cortical neurons may operate through CB1 and VGSCs, although this may not be the case in other parts of the CNS (Peters et al., 2010).

VGSC are directly responsible for the generation of action potentials, and therefore their strong modulation by AEA represents a powerful pathway by which AEA influences excitability. Phyto- and synthetic cannabinoid actions on neuronal excitability have become of increasing importance because of the widespread legalization of marijuana for medicinal and recreational use (Wilkinson et al., 2016). Our demonstration that VGSC inhibition by AEA occurs via CB1 provides an explanation for the antiepileptic and analgesic actions of exogenous cannabinoids (Benedetti et al., 2011; Holdcroft et al., 2006; Michalski et al., 2007; Perucca, 2017; Stockings et al., 2018). Enhanced understanding of these mechanisms may lead to the refinement of therapeutic cannabinoids that retain therapeutic effects but have fewer unwanted side effects.

### Compartmentalization of AEA signaling

Regional specificity is a fundamental attribute of GPCR signaling. In highly polarized cells such as neurons, it is possible that heterogeneity in GPCR signaling exists based on the proximity of key cellular machinery. AEA inhibition of VGSC is more tightly coupled in the soma. We found VGSC currents recorded from boutons and dendrites to be less sensitive to AEA (Figures 5 & 6). Furthermore, inhibition of terminal VGSC recordings was insensitive to CB1 deletion. Taken together these experiments indicate somatic CB1 is pharmacologically privileged in regulating VGSCs. This may be explained by variable expression of VGSC channel subtypes in distinct regions of the neuron that are selectively modulated by CB1 signaling cascade. Similar differential GPCR regulation of VGSC in the cortex has been reported. For instance, activation of the 5-HT_1A_ receptor inhibited VGSC current when measured at the axon initial segment (AIS), but not on the axonal trunk. This was reportedly due to separate VGSC populations, where NaV but not 1.6 was sensitive to serotonin (Yin et al., 2017). We observed complete VGSC current block after prolonged AEA application indicating VGSCs in the soma and AIS were both affected by AEA. We cannot exclude the possibility that VGSC subtypes in the dendrites and axons were less sensitive to AEA application. However, in dendrites it seems that the absence of CB1 (Figure 4) is the likely explanation for loss of AEA sensitivity. At terminals the answer may be more complicated, since CB1 receptors appear present in the minority of axons but deletion had no effect on the response to AEA, suggesting a lower efficacy CB1-independent pathway is involved. The basis for the reduced coupling between CB1 and VGSCs in the axons expressing CB1 is unclear. However, the dependence of the role for CB1 on the neuronal compartment reflects a separation of regulation between bouton and soma that has been observed for other proteins such as potassium channels (Li et al., 2020; Ritzau-Jost *et al*., 2021).

The enhanced efficacy of intracellular CB1 antagonists and the localization of CB1 to the intracellular compartment of the soma (Figures 3 and 4) in our experiments is consistent with other studies that show organelle-associated CB1 signaling in hippocampal neurons (Bénard *et al*., 2012; Hebert-Chatelain et al., 2016). Activation of the mitochondria-associated CB1 reduced respiratory chain activity and contributed to depolarization-induced suppression of inhibition linking energy metabolism and hippocampal neuroplasticity (Bénard *et al*., 2012; Hebert-Chatelain *et al*., 2016). Likewise, inhibition of VGSC currents by CB1 activation is a plausible mechanism to couple excitability and energy metabolism in the since action potential generation is energetically expensive (Hu et al., 2018). Presumably, if neocortical CB1 also couples to the respiratory chain, VGSC inhibition provides a mechanism by which AEA could decrease action potentials and ATP supply. Compartmentalization of CB1 could provide highly localized regulation of action potentials to optimize the trade-off between metabolic costs and conduction reliability (Hallermann et al., 2012).

In conclusion, our demonstration that CB1 activation causes VGSC inhibition with high efficacy in nearly all neocortical neurons describes an important role for AEA signaling in the CNS. The signaling pathway also provides a mechanism to explain the antiseizure and analgesic effects of cannabinoids.

## Methods

### Animals and genotyping

Control wild-type mice were obtained from a colony consisting of a stable strain of C57BL/6J x 129 × 1 mice. In addition, *Cnr1*^*+/+*^ and *Cnr1*^*-/-*^ mice were generated by breeding *Cnr1*^*+/-*^ pairs (Ledent et al., 1999) as described previously. CD1 was the background strain of the *Cnr1*^*+/-*^ mice. Breeding pairs were housed together on a 12-12 hour light-dark cycle with food and water available *ad libitum*. DNA extraction was performed using the Hot Shot method (Truett et al., 2000) with a 1-2 h boil. Primers used for CB1 PCRs were 5′-CATCATCACAGATTTCTATGTAC-3′ and 5′-GAGGTGCCAGGAGGGAACC-3′, to amplify a 366 bp band from the wild-type allele and 5′-GATCCAGAACATCAGGTAGG-3′ and 5′-AAGGAAGGGTGAGAACAGAG-3′, for a 521 bp band from the mutant CB1 allele.

### Neuronal Cell Culture

Neocortical neurons were isolated by whole brain dissection from P1-2 mouse pups of either sex as described previously (Martiszus et al., 2021). Briefly, mice were decapitated during general anesthesia with isoflurane, and the cerebral cortices were removed. Cortices were digested with trypsin and DNAse then dissociated with heat polished pipettes of varying diameters. Cells were then plated on Matrigel-coated (Corning, United States) glass coverslips and cultured in MEM supplemented with 5% FBS. To limit division of glial cells, cytosine arabinoside (4 µM) was added 48 hours later. All animal procedures were approved by the VA Portland Health Care System Institutional Animal Care and Use Committee in accordance with the U.S. Public Health Service Policy on Humane Care and Use of Laboratory Animals and the NIH Guide for the Care and Use of Laboratory Animals.

### Electrophysiological Recordings

Patch clamp experiments were performed using HEKA EPC10 USB amplifiers. Cells were visualized using a microscope (Olympus IX-70 or Scientifica) coupled to CCD-cameras (Andor iXon Ultra Camera or Scientifica FWCAM). Recordings were made from neocortical neurons after 7 to 60 days in culture. The cells were perfused with Tyrode solution that contained: (mM) 150 NaCl, 4 KCl, 10 HEPES, 10 glucose, 1.1 MgCl_2_, 1.1 CaCl_2_, pH 7.35 with NaOH. Voltage-clamp experiments used the following pipette solution:135mM CsMeSO_3_, 1.8 mM EGTA, 10 mM HEPES, 4 mM MgCl_2_, 0.2 mM CaCl_2_, 0.3 mM NaGTP, 5 mM NaATP, and 14 mM creatine phosphate (disodium salt), pH adjusted to 7.2 with TEA-OH. Current clamp recordings were made using the following pipette solution: 118 mM K-gluconate, 9 mM EGTA, 10 mM HEPES, 4 mM MgCl_2_, 1 mM CaCl_2_, 4 mM NaATP, 0.3 mM NaGTP, and 14 mM creatinine phosphate. Membrane potentials were adjusted for the liquid junction potentials. Series resistance was compensated between 60-80%. Excitatory and inhibitory transmission was blocked by adding the following blockers to the extracellular solution: 10 µM CNQX, 50 µM APV, and 10 µM Gabazine (Abcam, United Kingdom).

### Data Acquisition and Analysis

Current and voltage traces were acquired using PatchMaster (HEKA, Germany). Signals were digitized at 50 kHz and filtered at 2.9 or 5 kHz. Leak current subtraction employed p/n protocols, with currents elicited using voltage step sizes selected to minimize artefacts. Analysis was performed using custom applications developed for Igor Pro (Wavemetrics, Lake Oswego, OR). Time course experiments were normalized to the average baseline currents recorded 50-100 s before the drug application. Data values were reported as mean (± SEM) or median, if not normally distributed. Statistical significance was determined with appropriate parametric or non-parametric tests (Excel, Graphpad prism or Igor Pro).

### Solutions

Solutions were gravity-fed via a 1.2 mm OD glass capillary tube located 2 mm from the target neuron. Most reagents were obtained from Sigma-Aldrich (Darmstadt, Germany). Anandamide (Abcam, United Kingdom) was acquired as a solution in purged ethanol (final concentration 0.07%). Solutions were switched manually using a low dead volume manifold upstream of the glass capillary. Vehicle control experiments utilized the same concentrations of ethanol and DMSO that were used in the paired experiment. For a full list of solutions see the Reagent Table.

### Immunocytofluorescence

Cells were fixed by placing the coverslip in 4% (v/v) paraformaldehyde for 10 min. After washing three times with phosphate-buffered saline (PBS), cells were blocked and permeabilized with PBS containing 1% bovine serum albumin (BSA), 5% normal goat serum (NGS) and 0.2% saponin at room temperature (30 min.), then incubated overnight at 4°C with 1:1000 mouse anti-Synaptophysin1 and rabbit anti-CB1 monoclonal antibodies (Synaptic Systems) diluted in the blocking solution. The next day cells were washed three times with PBS, blocked again for 30 min., and incubated with goat anti-mouse AlexaFluor 488 (1:1000) and goat anti-rabbit AlexaFluor 594 diluted in blocking solution for 60 min. Thereafter they were again washed three times (5 min.) with PBS. Coverslips were then mounted in Flouromount G reagent (SouthernBiolabs).

## Acknowledgements

We are grateful to Drs. David Farrens, James Frank, Salil Rajayer, and Eric Schnell for helpful discussions. The authors declare no competing financial interests. The contents do not represent the views of the U.S. Department of Veterans Affairs or the United States Government. This work was supported by grants awarded by NIGMS (GM134110) and U.S. Department of Veterans Affairs (BX002547) to SMS and NHLBI (T32HL083808) and NINDS (F31NS095463) to GBM. We thank Drs. Catherine Ledent and Ken Mackie for their kind provision of the CB1 knockout mice.

## Notes

### Competing Interest Statement

The authors have declared no competing interest.

