## Supplemental Figure 1 for "Somatic and terminal CB1 receptors are differentially coupled to voltage-gated sodium channels in neocortical neurons"

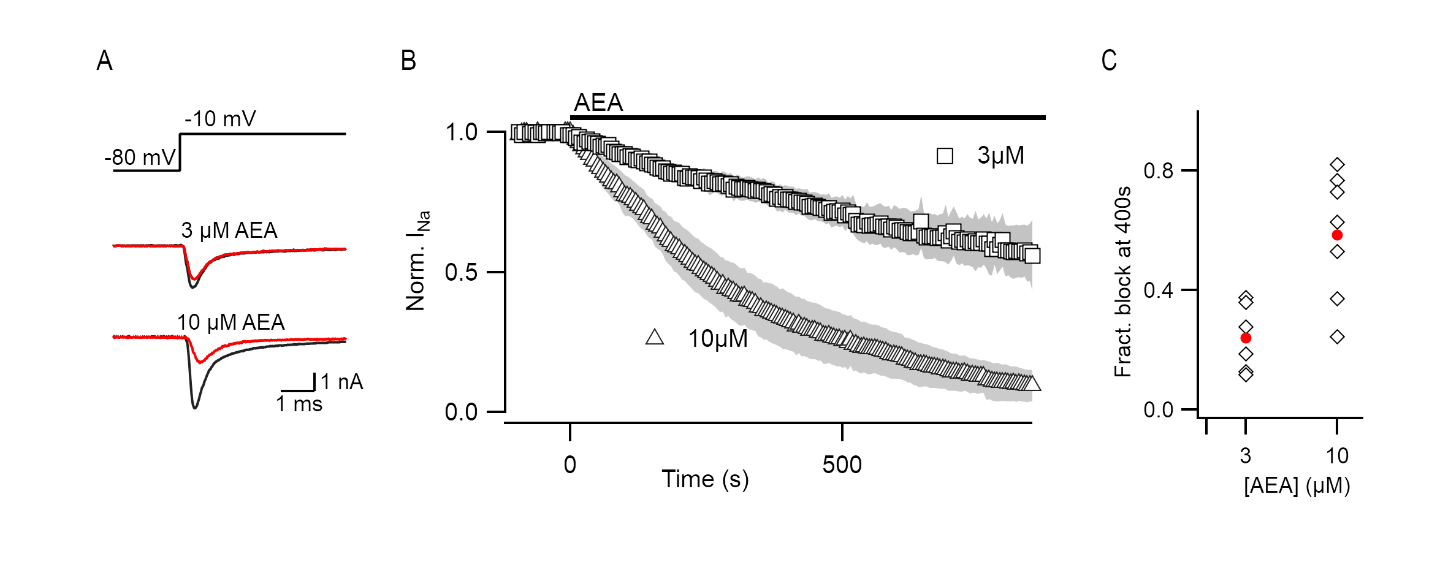

Supplement 1.

A) Exemplar VGSC traces of neocortical neurons before (black) and 400 seconds after (red) perfusion of either 3 or 10 µM AEA.
B) Time course of VGSC current amplitude prior to and during perfusion of either 3 (squares, n = 6) or 10 (triangles, n = 7) µM. AEA
C) Fractional block of VGSC current after 400 seconds of AEA application of 3 or 10 µM AEA.
